# SARS-CoV-2 growth, furin-cleavage-site adaptation and neutralization using serum from acutely infected, hospitalized COVID-19 patients

**DOI:** 10.1101/2020.06.19.154930

**Authors:** William B. Klimstra, Natasha L. Tilston-Lunel, Sham Nambulli, James Boslett, Cynthia M. McMillen, Theron Gilliland, Matthew D. Dunn, Chengqun Sun, Sarah E Wheeler, Alan Wells, Amy L. Hartman, Anita K. McElroy, Douglas S. Reed, Linda J. Rennick, W. Paul Duprex

## Abstract

SARS-CoV-2, the causative agent of COVID-19, emerged at the end of 2019 and by mid-June 2020, the virus has spread to at least 215 countries, caused more than 8,000,000 confirmed infections and over 450,000 deaths, and overwhelmed healthcare systems worldwide. Like SARS-CoV, which emerged in 2002 and caused a similar disease, SARS-CoV-2 is a betacoronavirus. Both viruses use human angiotensin-converting enzyme 2 (hACE2) as a receptor to enter cells. However, the SARS-CoV-2 spike (S) glycoprotein has a novel insertion that generates a putative furin cleavage signal and this has been postulated to expand the host range. Two low passage (P) strains of SARS-CoV-2 (Wash1: P4 and Munich: P1) were cultured twice in Vero-E6 cells and characterized virologically. Sanger and MinION sequencing demonstrated significant deletions in the furin cleavage signal of Wash1: P6 and minor variants in the Munich: P3 strain. Cleavage of the S glycoprotein in SARS-CoV-2-infected Vero-E6 cell lysates was inefficient even when an intact furin cleavage signal was present. Indirect immunofluorescence demonstrated the S glycoprotein reached the cell surface. Since the S protein is a major antigenic target for the development of neutralizing antibodies we investigated the development of neutralizing antibody titers in serial serum samples obtained from COVID-19 human patients. These were comparable regardless of the presence of an intact or deleted furin cleavage signal. These studies illustrate the need to characterize virus stocks meticulously prior to performing either *in vitro* or *in vivo* pathogenesis studies.

## Introduction

On December 31, 2019 Chinese authorities notified the World Health Organization (WHO) of a pneumonia of unknown etiology (1). As of June 18, 2020, the disease, subsequently named coronavirus disease 2019 (COVID-19) by the WHO (ref https://www.who.int/docs/default-source/coronaviruse/situation-reports/20200211-sitrep-22-ncov.pdf), had spread to 215 countries with 8,385,440 confirmed cases and 450,686 confirmed deaths worldwide (https://www.who.int/emergencies/diseases/novel-coronavirus-2019, accessed June 19, 2020. Severe acute respiratory syndrome coronavirus 2 (SARS-CoV-2) (2) was identified as the causative agent in January 2020 (3). SARS-CoV-2 is an enveloped, non-segmented, single-stranded, positive-sense, RNA virus, and has been classified as a betacoronavirus alongside severe acute respiratory syndrome coronavirus (SARS-CoV), which emerged in China in 2002 (4, 5), and Middle East respiratory syndrome coronavirus (MERS-CoV) which emerged in the Arabian Peninsula in 2012 (6). SARS-CoV and MERS-CoV are most likely bat viruses which infect humans via intermediate hosts (7-9). Although the reservoir species remains unidentified, SARS-CoV-2 is also thought to be a zoonotic bat virus (10).

The SARS-CoV-2 genome encodes structural proteins, including the nucleo- (N), membrane (M), envelope (E) and spike (S) proteins, non-structural proteins and accessory proteins (3). The multifunctional coronavirus N protein primarily encapsidates the viral (v) RNA and plays a role in virus replication, transcription and assembly, as well as modulating the host cell environment (11). The coronavirus E protein is an integral membrane protein that forms viroporins, complexes essential for virus assembly and release (12). The transmembrane glycoprotein, coronavirus M protein, is located in the viral envelope and is required for egress (13). The S protein of coronaviruses is a type I fusion protein involved in attachment to human angiotensin-converting enzyme 2 (hACE2) at the cell membrane (14) and subsequent virus-to-cell fusion which in turn liberates the vRNA into the cytoplasm (15). To accomplish this, most coronavirus S proteins undergo host cell-mediated cleavage to generate the S1 protein, responsible for receptor binding and the S2 protein, responsible for virus-to-cell fusion (16). The S2 polypeptide is further processed at the S2’ site to expose the fusion peptide (17, 18). Coronavirus S protein processing and entry strategies vary. For example, the S protein of the prototypic coronavirus mouse hepatitis virus (MHV), is cleaved during biosynthesis in the infected cell whereas the S protein of SARS-CoV remains uncleaved until receptor binding and entry of progeny virus during subsequent infections (19).

Several proteases process coronavirus S proteins depending on where and when this takes place. Proprotein convertases, extracellular proteases, cell surface proteases or lysosomal proteases can process the protein depending on whether spikes are cleaved during virus packaging (e.g. furin), after virus release (e.g. elastase), after virus attachment to the host cell (e.g. TMPRSS2), or during virus entry by endocytosis (e.g. cathepsin L), respectively (20). Sequence analysis of SARS-CoV-2 S protein from human cases shows an insertion at the S1/S2 boundary *versus* the proposed bat reservoir virus. This generates a novel, putative, furin cleavage signal not seen in other clade members (21). It was subsequently shown that prior furin cleavage enhanced entry of a pseudovirus containing the SARS-CoV-2 S protein into different cell lines expressing hACE2 (22, 23). The furin cleavage site is also required for SARS-CoV-2 infection of human lung cells (24).

The S protein, along with the N and M proteins, is also a major antigenic target. A study of T cell responses to SARS-CoV-2 found that CD4^+^ cells were directed predominantly against S, N and M proteins and CD8^+^ cells were directed predominantly against S and M proteins, with significant reactivity also against N protein, along with some non-structural proteins (25). Using an N-protein ELISA, IgM and IgA antibodies have been detected as early as three days post symptom onset (d.p.s.o.) with IgG antibodies detected as early as ten d.p.s.o. (26). The S protein is the major target for neutralizing antibodies (27, 28) and neutralizing antibody titers were reported to be higher in older COVID-19 patients (28). Two linear B-cell epitopes have been reported on the S protein and it was shown that antibodies recognizing these epitopes accounted for a high proportion of the anti-S neutralizing antibody response (29).

In this study we aimed to characterize SARS-CoV-2 isolates from two sources virologically and immunologically and determine the impact passage in Vero-E6 cells had on the genetic sequence of the furin cleavage site. We examined the effect of virus passage on stability of the furin cleavage signal at the S1/S2 boundary, and subsequently investigated if there are differences in neutralizing antibody titers for viruses that have different sequences in this region. Neutralization activity was assessed using serial serum samples from acutely infected COVID-19 patients. Such analyses are critical for standardization of virus strains prior to generation of cDNA clones, examination of *in vitro* cell interaction characteristics, pathogenesis studies and identification of appropriate challenge strains for vaccine and therapeutic efficacy studies.

## Methods

Virus growth and assays were performed at Biosafety Level 3 (BSL-3) conditions in the Regional Biocontainment Laboratory (RBL) in the Center for Vaccine Research, at the University of Pittsburgh. All infectious material was inactivated using University of Pittsburgh Biohazards/Biosafety committee-approved protocols prior to removal from BSL-3 to allow assays to be conducted at lower containment.

### Cells

Vero 76 cells, Clone E6, were purchased from ATCC and were grown in DMEM (Corning) supplemented with 10% (v/v) FBS (Atlanta Biologicals), 1% (v/v) L-glutamine (Corning) and 1% (v/v) penicillin-streptomycin (pen-strep; Corning).

### Serum samples

De-identified, serum samples from three acutely infected COVID-19 (PCR-confirmed) patients treated at University of Pittsburgh Medical Center (UPMC) were received, aliquoted at BSL-2+ (Table 1). These specimens were obtained as excess pathological specimens under IRB approved protocols.

**Table 1.**
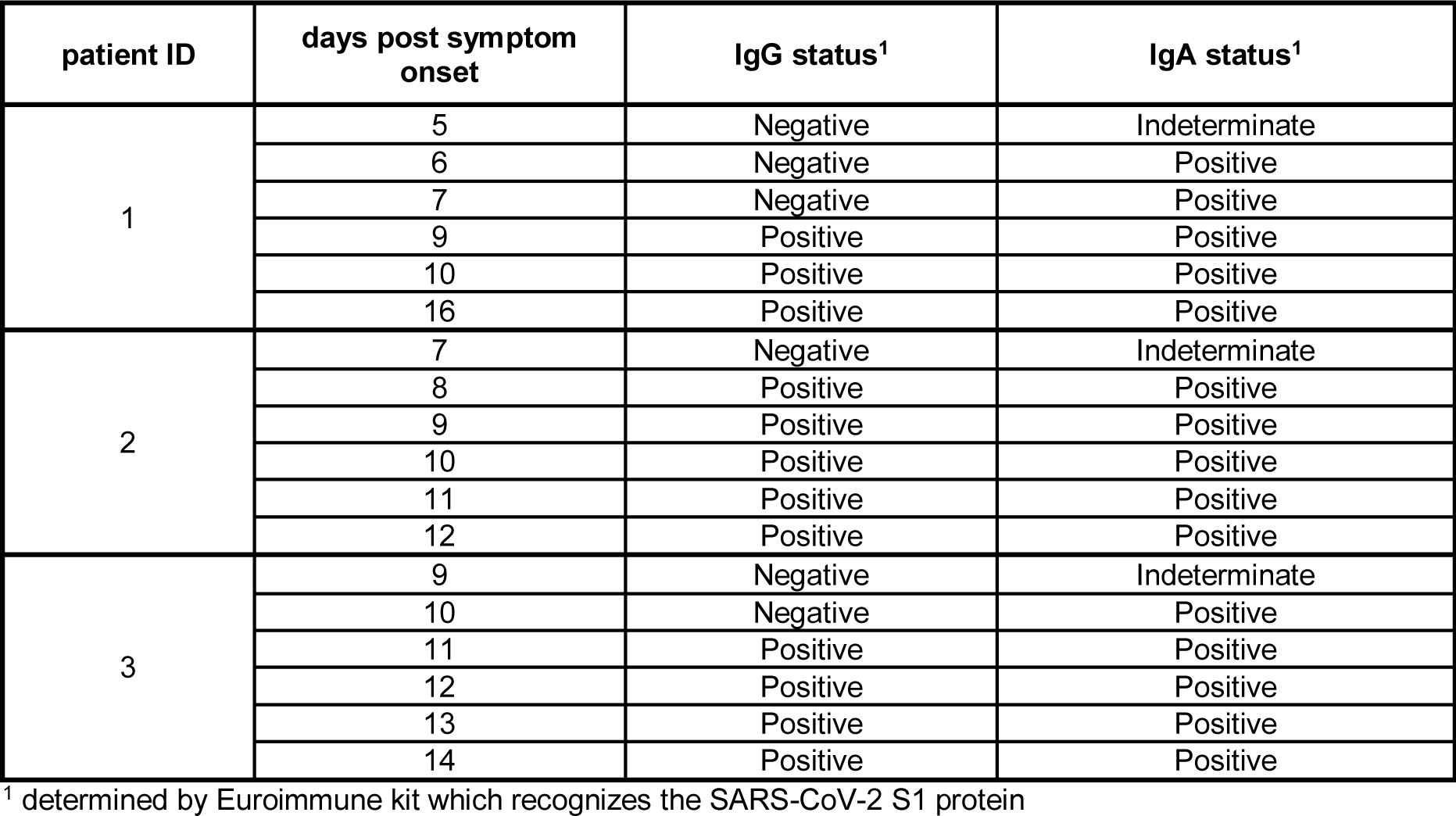
Serum samples used for plaque reduction neutralization and indirect immunofluorescence assays

| patient ID | days post symptom onset | IgG status<sup>1</sup> | IgA status<sup>1</sup> |
| --- | --- | --- | --- |
| 1 | 5 | Negative | Indeterminate |
|  | 6 | Negative | Positive |
|  | 7 | Negative | Positive |
|  | 9 | Positive | Positive |
|  | 10 | Positive | Positive |
|  | 16 | Positive | Positive |
| 2 | 7 | Negative | Indeterminate |
|  | 8 | Positive | Positive |
|  | 9 | Positive | Positive |
|  | 10 | Positive | Positive |
|  | 11 | Positive | Positive |
|  | 12 | Positive | Positive |
| 3 | 9 | Negative | Indeterminate |
|  | 10 | Negative | Positive |
|  | 11 | Positive | Positive |
|  | 12 | Positive | Positive |
|  | 13 | Positive | Positive |
|  | 14 | Positive | Positive |

<sup>1</sup> determined by Euroimmune kit which recognizes the SARS-CoV-2 S1 protein

### Virus stock production and passage

Passage 4 CDC/2019n-CoV/USA_WA1/2020 (Wash1) or P1 SARS-CoV-2/München- 1.1/2020/929 (Munich) virus was diluted to 500 µl in 15 ml with virus dilution medium (Opti-MEM, Gibco) supplemented with 2% (v/v) fetal bovine serum (FBS) and added to a confluent monolayer of Vero-E6 cells in an expanded-surface 1900 cm^2^ roller bottle (CELLTREAT Scientific Products). After 1 hour of incubation at 37 °C, 5% (v/v) CO_2_, 45 ml of virus growth medium (DMEM supplemented with 10% (v/v) FBS, 1% (v/v) L-glutamine and 1% (v/v) pen-strep) was added and cells were incubated under the same conditions for 96 hours. Virus containing-supernatant was collected and clarified by centrifugation at 3500 rpm for 30 minutes at 4°C. The cleared virus supernatant was aliquoted and stored at -80 °C.

To generate large volume virus stocks, 60 ml of Wash1: P5 or Munich: P2 virus was thawed, diluted to 125 ml with virus dilution medium and added to confluent monolayers of Vero-E6 cells in twenty-five T175 flasks (Nunc; 5 ml/flask). After 1 hour of incubation at 37 °C, 5% (v/v) CO_2_, 20 ml/flask of virus growth medium was added and incubation was continued for 66-72 hours until cytopathic effect was observed. Virus containing-supernatant was collected and clarified by centrifugation at 3500 rpm for 30 minutes at 4 °C. The cleared virus supernatant was aliquoted and stored at -80 °C.

### Plaque assay

Ten-fold serial dilutions of virus stock were prepared in virus dilution medium and were added, in duplicate, to confluent Vero-E6 monolayers in six-well plates (Corning; 200 µl/well). After 1 hour of incubation at 37 °C, 5% CO_2_, 2 ml/well of virus growth medium containing 0.1% (w/v) immunodiffusion agarose (MP Biomedicals) was added and incubation was continued for 96 hours. Plates were fixed with 2 ml/well formaldehyde (37% (w/v) formaldehyde stabilized with 10- 15% (v/v) methanol; Fisher Scientific) for 15 minutes at room temperature. Agarose and fixative were discarded and 1 ml/well 1% (w/v) crystal violet in 10% (v/v) methanol (both Fisher Scientific) was added. Plates were stored at room temperature for 20 minutes and then rinsed thoroughly with water. Plaques were then enumerated.

### RNA isolation

400 µl of Tri-Reagent (ThermoFisher) was added to 100 µl of virus stock and incubated for 10 minutes at room temperature. 5 μl of polyacryl carrier (Molecular Research Center) was added to the specimen/Tri-Reagent mixture, and incubated at room temperature for 30 seconds. Next, 200 μl of chloroform was added to the sample which was inverted vigorously for 30 seconds before centrifugation at 12,000 rpm for 15 minutes at 4 °C. The aqueous phase was collected, mixed with an equal volume of 100% isopropanol and incubated for 10 minutes at room temperature before centrifugation at 12,000 rpm for 10 minutes at 4 °C to pellet the RNA. The supernatant was removed and the RNA pellet was washed with 1 ml of 70% (v/v) ethanol before centrifugation at 12,000 rpm for 5 minutes. The wash and centrifugation step was repeated after which the supernatant was removed and the RNA pellet was allowed to air dry for 5 minutes before suspending in 40 μl of nuclease-free water. The RNA was stored at -80 °C until reverse transcription (RT) quantitative polymerase chain reaction (RT-qPCR) and sequence analyses.

### Quantitative RT-qPCR

For quantitation of genomic vRNA, one-step RT-qPCR was performed using the 4X Reliance One-Step Multiplex RT-qPCR Supermix (BioRad), following the manufacturer’s instructions. Primers and probe targeting the N gene sequence were as designed and optimized by the Centers for Disease Control and Prevention and sequences (Table 2). The working concentrations for the forward and reverse primers and probe were 20 μM, 20 μM, and 5 μM, respectively. Reverse

**Table 2:**
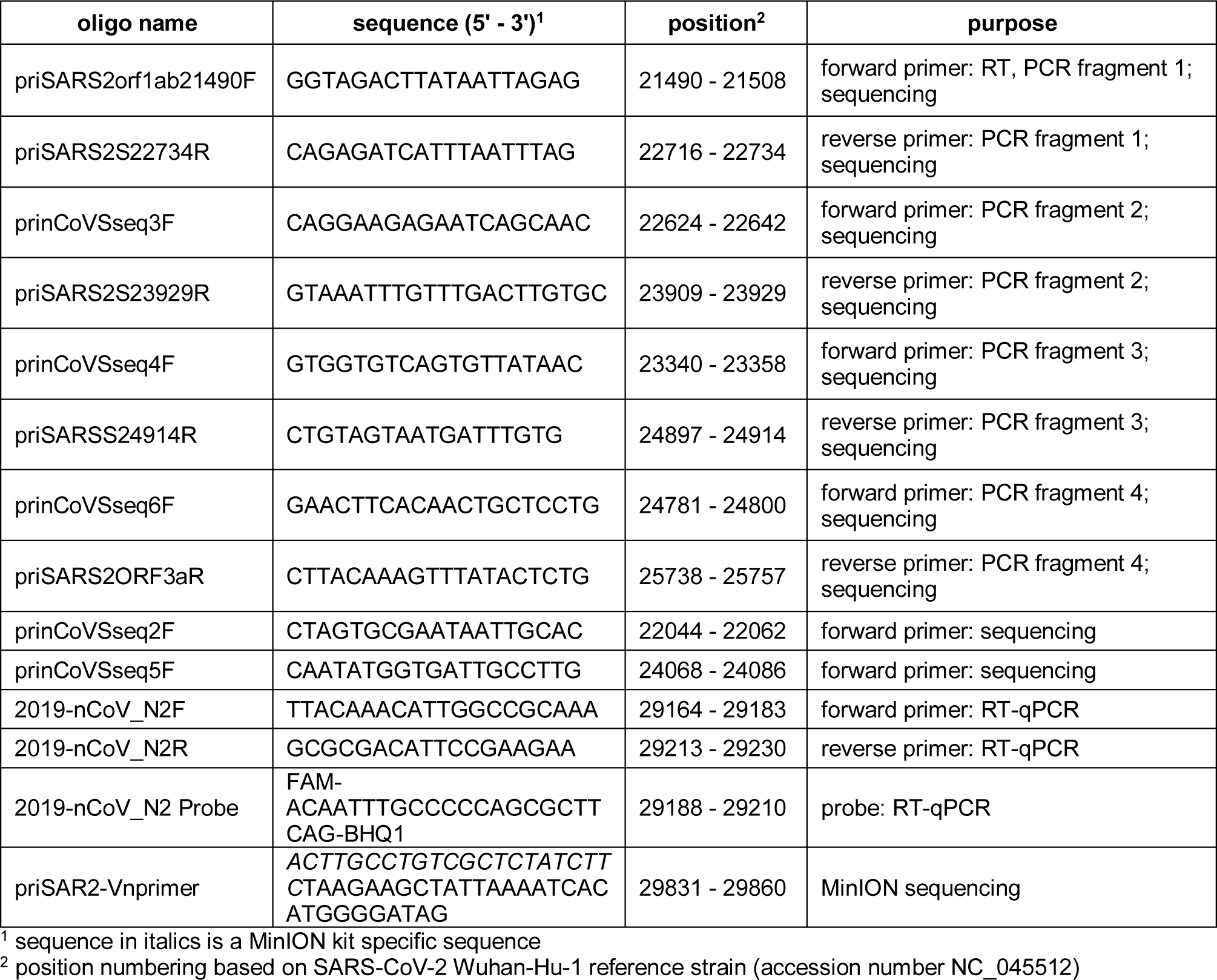
DNA oligos used for reverse transcription, polymerase chain reaction and sequencing

transcription occurred at 50 °C for 10 minutes, followed by a polymerase activation step at 95 °C for 10 minutes, and a cycling PCR amplification (40 cycles) comprising 95 °C for 10 seconds and 60 °C for 30 seconds. Quantitation of virus genome copies was determined by comparing the cycle threshold (CT) values from the unknown samples to CT values from a positive-sense SARS-CoV-2 vRNA standard curve generated from 1:10 serial dilutions of template which was synthesized in-house by *in vitro* transcription using the mMessage mMachine T7 kit (Ambion) and following the manufacturer’s instructions. The limit of detection (LOD) was 23.2 genome copies.

### Sanger sequencing

Reverse transcription-polymerase chain reaction (RT-PCR) was performed on virion RNA from virus stock passages using a range of primers (Table 2). Reactions were performed with a SuperScript III First-Strand Synthesis System (Invitrogen) using a virus-specific forward primer that bound in orf1ab. The cDNAs encoding S gene sequences were amplified by PCR, either in their entirety or as four overlapping fragments, were carried out using Phusion High-Fidelity DNA Polymerase (New England Biolabs). Amplicons were agarose gel-extracted and purified using a QIAquick gel extraction kit (Qiagen) before Sanger sequencing (Genewiz, New Jersey).

### Oxford Nanopore (MinION) cDNA-PCR sequencing

SARS-CoV-2 transcripts from Wash1: P6 and Munich: P3 virus stocks were sequenced on the MinION device (MinION Mk1B, MIN-101B) using ONT Ligation Sequencing Kit 1D (SQK- LSK108). Modifications were made to the RT-PCR stage of library preparation according to manufacturer’s recommendations for sequencing Kit SQK-LSK109. In brief, 50 ng of RNA, quantified using a Qubit Fluorometer (Qubit 4, Life Technologies; Qubit RNA HS Assay Kit), was mixed with an oligo (2 µM, priSARS2-Vnprimer, Table 2) designed to bind upstream of the polyA-tail at the 3’ end of the genome and dNTPs (10 mM, NEB), and incubated at 65 °C for 5 minutes. RT buffer (Maxima H Minus Reverse Transcriptase, ThermoFisher), RNase OUT (Life Technologies) and strand-switching primer (10 µM, Kit SQK-LSK108) were added to the RNA-primer mix and incubated at 42 °C for 2 minutes. 200 U of Maxima H Minus Reverse Transcriptase was then added and first-strand cDNA synthesis performed (total volume 20 µl; RT and strand-switching at 42 °C for 90 minutes, heat inactivation at 85 °C for 5 minutes). Four PCR reactions were set-up using 5 µl of the cDNA template, 2X LongAmp *Taq* DNA polymerase, 2X Master Mix (NEB) and cDNA primer (cPRM, Kit SQK-LSK108). PCR conditions were set as initial denaturation for 30 seconds at 95°C, followed by 18 cycles of denaturation for 15 seconds at 95°C, annealing for 15 seconds at 62°C and extension for 25 minutes at 65°C, followed by final extension for 10 minutes at 65°C. Exonuclease I (20 U, NEB) was added to each PCR reaction and incubated at 37 °C for 15 minutes, followed by heat inactivation for 15 minutes. PCR reactions were pooled and purified with Agencourt AMPure XP beads (Beckman Coulter). Purified cDNA was eluted in 21 µl Rapid Annealing Buffer (RAB, Kit SQK-LSK108) and quantified using a Qubit Fluorometer (Qubit 4, Life Technologies; Qubit (ds)DNA HS Assay Kit). 5 µl of cDNA Adapter Mix (cAMX, Kit SQK-LSK108) was added to the cDNA library and incubated for 5 minutes at room temperature on a rotator, followed by incubation with Adapter Bead Binding buffer (ABB, Kit SQK-LSK108). The cDNA libraries were purified with Agencourt AMPure XP beads and eluted in 12 µl Elution Buffer (ELB, Kit SQK-LSK108). Each cDNA library was loaded individually onto separate SpotON Flow Cells (Version R9; FLO-MIN 106) and run using the default script for sequencing Kit SQK-LSK108 via MinKNOW. Live base calling, FAST5 and FASTQ outputs were turned on and sequencing occurred over 18 hours.

FASTQ files were imported into CLC Genomics Workbench 20.04 (Oxford Nanopore High-Throughput Sequencing Import). Passed reads were then aligned to SARS-CoV-2 Wash1 genome (accession number: MN985325) using the ‘Map Long Reads to Reference’ tool available through the ‘Long Read Support’ plugin (Beta). Consensus sequences were extracted and used for analysis against the published Wash1 genome sequence.

### Indirect immunofluorescence assay

Confluent monolayers of Vero-E6 cells in 35 mm-glass bottom dishes (Ibidi) were infected with P3 Munich virus at a multiplicity of infection (MOI) of 0.01. After 24 hours of incubation at 37 °C, 5% CO_2_, the supernatant was discarded, the cell monolayer was washed with phosphate buffered saline (PBS) and 4% (w/v) paraformaldehyde in PBS was added to fix. After 20 minutes of incubation at room temperature, the fixative was removed and the monolayer was washed twice with PBS. The cells were permeabilized with 0.1% (v/v) Triton X (Sigma) in PBS by incubation at 37 °C for 30 minutes. Primary antibody (rabbit anti-SARS-CoV-2 S monoclonal antibody, 40150- R007, Sino Biological, rabbit anti-SARS-CoV-2 N monoclonal antibody, 40143-R019, Sino Biological, or the day 14 serum from patient 3 (Table 1), diluted 1:500 in PBS containing 0.1% (v/v) Triton X was added and incubated at 37 °C for 1 hour. After washing the cells three times with PBS, secondary antibody (chicken anti-rabbit Alexa Fluor-488, Invitrogen, or goat anti-human Alexa Fluor-488, Invitrogen) diluted 1:400 in PBS containing 0.1% (v/v) Triton X was added and incubated at 37 °C for 1 hour. After washing the cells three times with PBS, 4′,6-diamidino-2- phenylindole (DAPI) nuclear stain (300 nM DAPI stain solution; Invitrogen) was added and incubated in the dark for 10 minutes at room temperature. For some assays, after washing the cells with PBS, Alexa Fluor-568 phalloidin (Invitrogen) was added to counterstain the actin cytoskeleton. After incubation for 30 minutes at room temperature, the cells were washed twice with PBS, and stored (protected from light) in PBS at 4 °C until imaging. Fluorescence was observed with a UV microscope (Leica) and photomicrographs were obtained using a camera (Leica) and LAS X software (Leica).

### Immunoplaque assay

Confluent monolayers of Vero-E6 cells in a 12-well plate were infected with 150 µl of virus inoculum from a 10-fold serial dilution prepared in Opti-MEM I. Cells were incubated for 1 hour at 37 °C and overlaid with 0.6% (v/v) Avicel (FMC Biopolymer) in 2X MEM (10X MEM, no glutamine, Gibco) supplemented with 2% (v/v/) FBS. Cells were incubated at 37 °C, 5% (v/v) CO_2_ for 2 days before fixing with 4% (w/v) paraformaldehyde in PBS for 30 minutes at room temperature. Fixative was removed and cells were washed with water (1 ml). Permeabilization buffer (0.5% Triton X-100, 20 mM sucrose in PBS; 1 ml) was added and incubated for 30 minutes at room temperature. Permeabilization buffer was removed and cells were washed with wash buffer (0.1% Tween-20 in PBS; 1 ml). Cells were incubated with 200 µl primary antibody (rabbit anti-SARS-CoV-2 N, Sino Biological, 40143-R019) diluted 1:1000 in blocking buffer (4% dried milk/0.1% Tween-20 in PBS) for 1 hour at room temperature. Primary antibody was removed and cells were washed three times with wash buffer. Cells were incubated with 200 µl secondary antibody (goat anti-rabbit HRP, Abcam, ab6721) diluted 1:1000 in blocking buffer for 1 hour at room temperature. Secondary antibody was removed and cells were washed three times with wash buffer. Cells were incubated with 200 µl KPL TrueBlue Peroxidase Substrate solution (Sera Care, 5510-0050) on a rocker for 20 minutes at room temperature. The substrate was removed and the cells were washed once with water (1 ml) to stop the reaction. Plates were digitalized using a light box and 13-megapixel Nikon digital camera.

### Western blot

Confluent monolayers of Vero-E6 cells in 6-well plates were infected with Wash1 (P5 or P6) or Munich (P2 or P3) virus at an MOI of 0.01 or transfected with a pCAGGS vector expressing a codon-optimized (GeneScript) S protein (synthesized by GeneScript) possessing the amino acid sequence of the SARS-CoV-2 Wuhan-Hu-1 isolate (accession number MN908947). After 48 hours of incubation at 37 °C, 5% (v/v) CO_2_, the supernatant was discarded and 200 µl of RIPA buffer (50 mM TRIS-HCl pH 7.4, 150 mM NaCl, 1% (v/v) NP-40, 0.5% (v/v) sodium deoxycholate, 0.11 mM EDTA, 0.11 mM EGTA, and 1.1 mM DTT, all Sigma with Protease Inhibitor Cocktail Tablets, REF 04693124001, Roche) was added. After incubation at room temperature for 5 minutes, the monolayer was scraped into solution using the rubber tip of a plunger from a 1 ml syringe (Fisher Scientific) and stored at -80 °C until use.

Cell lysates in 1X RIPA buffer were mixed with an equal volume of 2X Laemmli sample buffer (Bio-RAD, containing 50 µl/ml 2-mercaptoethanol) followed by boiling for 5 minutes. Equal amounts of protein were loaded into the wells of 8% (w/v) SDS-PAGE gels which were run in running buffer (25 mM TRIS-HCl, 192 mM glycine, 0.1% SDS pH 8.3) for 1–2 hours at 100 V. Proteins were transferred to polyvinylidene difluoride (PVDF) membranes in 1X transfer buffer (wet) for 3 hours at 0.22 mA. The membranes were blocked with 5% (w/v) non-fat milk in TBS-Tween-20 (TBST) buffer (Thermo Scientific) for 1 hour at room temperature. The membranes were incubated with rabbit anti-SARS coronavirus spike protein polyclonal antibody (Invitrogen, PA1-41513) diluted 1:10,000, or rabbit anti-SARS-CoV-2 N monoclonal antibody (Sino Biological, 40143-R019) diluted 1:15,000, in TBST buffer containing 1% (w/v) non-fat milk overnight at 4 °C. The antibodies were removed and the membranes washed five times for 10 minutes with TBST buffer. The membranes were incubated with HRP-goat anti-rabbit IgG (Invitrogen, REF656120) diluted 1:20,000 in TBST buffer containing 1% (w/v) non-fat milk for 1 hour at room temperature. The antibody was removed and the membranes were washed five times for 10 minutes with TBST buffer. The membranes were incubated with a working solution of enhanced chemiluminescence ECL western blotting substrate (Pierce) for 5 minutes at room temperature. Excess substrate was removed and the membrane was exposed to X-ray film.

### Plaque reduction neutralization assay

Human serum samples (Table 1) were diluted in a 2-fold series from 1:50 to 1:1600 in Opti-MEM I. Each serum dilution (100 µl) was mixed with 100 µl of Munich: P3 or Wash1: P6 virus containing 50 plaque forming units (P.F.U.) of virus in Opti-MEM I. The serum-virus mixes (200 µl total) were incubated at 37 °C for 1 hour after which they were added dropwise onto confluent Vero-E6 cell monolayers in 6-well plates. After incubation at 37 °C, 5% (v/v) CO_2_ for 1 hour, 2 ml of 0.1% (w/v) immunodiffusion agarose in DMEM supplemented with 10% (v/v) FBS and 1X pen-strep was added to each well. After incubation at 37 °C, 5% (v/v) CO_2_ for 72 hours, the agarose overlay was removed and the cell monolayer was fixed with 1 ml/well formaldehyde (37% (w/v) formaldehyde stabilized with 10-15% (v/v) methanol) for 20 minutes at room temperature. Fixative was discarded and 1 ml/well 1% (w/v) crystal violet in 10% (v/v) methanol was added. Plates were incubated at room temperature for 20 minutes and rinsed thoroughly with water. Plaques were then enumerated and the 80% plaque reduction neutralization titer (PRNT_80_) was calculated. Controls, using validated SARS-CoV-2 antibody-negative human serum control and an uninfected cells control, were performed to ensure virus neutralizing was specific.

## Results

### Virus passage and characterization

Two viruses isolated from clinical samples were obtained. CDC/2019n-CoV/USA_WA1/2020 (Wash1) was isolated from the first COVID-19 case in Washington state, United States and was received after four passages in Vero CCL-81 cells (P4) (30). A second virus, SARS-CoV- 2/München-1.1/2020/929 (Munich) was isolated from a hospitalized patient in Munich, Germany and was received directly after the initial isolation in Vero-E6 cells (P1) (31). We further passaged both viruses twice in Vero-E6 cells to obtain large volume working stocks at P6 and P3 respectively since these cells are known to support efficient production (30). An agarose overlay plaque assay was developed and each virus stock was titrated by plaque assay on Vero E6 cells. Titers of 3.55×10^6^ P.F.U./ml and 3.25×10^5^ P.F.U./ml were obtained for Wash1: P6 and Munich: P3, respectively. Passage resulted in an apparent increase in the overall plaque size for both viruses (Fig. 1A and B). An immunoplaque assay was developed to ensure a) that the virus was SARS-CoV-2 and b) that all plaques could be identified. SARS-CoV-2 antigens were detected using an anti-N polyclonal antibody (Fig. 1C). Variation in plaque size was observed for both strains and the immunoplaque assay was useful in determining accurate viral titers since it was discriminatory between infected and uninfected cells.

**Figure 1.**
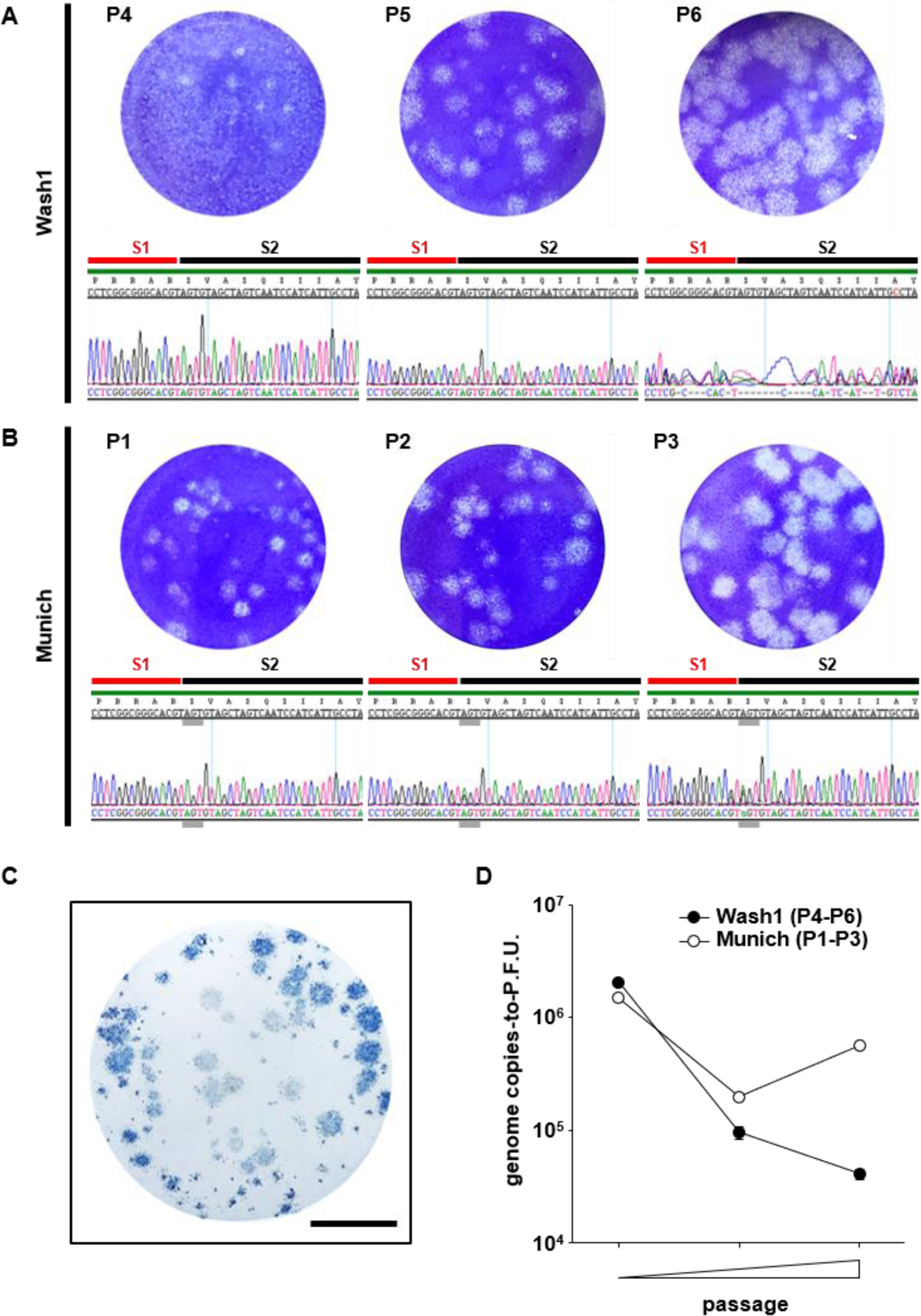
Virus growth and characterization. (A and B) Representative images showing plaques generated by infection of Vero-E6 cells with different passages of the Wash1 (A) and Munich (B) virus stocks. Chromatograms under each plaque image show the nucleotide sequence coding for the furin cleavage signal (RRAR) region of the S protein for each virus passage. Red and black lines indicate the putative S1 and S2 subunits, respectively, generated after cleavage. Grey lines indicate the nucleotide triplet coding for S 686. (C) Immunodetection of plaques (blue), using an anti-SARS-CoV-2 N antibody and HRP-conjugated secondary, confirming that plaques produced in Vero-E6 monolayers represent SARS-CoV-2 infection. The scale bar represents 1 cm. (D) Change in genome copies-to-P.F.U. ratio for virus stocks over two passages in Vero-E6 cells (Wash1: P4-P6; Munich: P1-P3). RNA was extracted from cell free supernatant virus stock and genome copies were enumerated by RT-qPCR. Virus stock titers were determined by plaque assay. Error bars represent one standard error of the mean.

Virus passage *in vitro*, particularly in Vero cells, can alter the virus particle-to-P.F.U. ratio as well as plaque sizes as the virus adapts to the cells used for culture (32, 33). We examined the genome copies- (as a proxy for particles) to-P.F.U. ratio for each virus at each passage (Fig. 1D). For the Wash1 virus the ratio decreased >10-fold from 2×10^6^ to 4×10^4^ between P4 and P6. However, for the Munich virus, the ratio dropped from 1.5×10^6^ at P1 to 2×10^5^ at P2 but then partially recovered to 5.5×10^5^ on the next passage.

Amplicons from Wash1: P4, P5 and P6 and Munich: P1, P2 and P3 stocks were examined by Sanger consensus sequencing across the S1 (red line)/S2 (black line) junction (Fig. 1A and B). Overlapping peaks were absent in the electrochromatograms for P1 and P4 but appeared in subsequent passages. The Wash1: P4 virus stock originally received from CDC and a P5 virus stock had S protein sequences that exactly matched the published sequence in this region. Sequencing using forward and reverse primers demonstrated significant heterogeneity around the S1/S2 junction in the Wash1: P6 passaged virus (Fig. 1A, right hand panel). Changes were undetectable in the Munich: P1 stock but one position demonstrated heterogeneity as a minor peak on the chromatograms for a P2 stock and was enriched on the P3 electrochromatograms (Fig. 1B, grey line). Therefore, Nanopore MinION long read sequencing was used to examine the genetic diversity in both the Wash1: P6 and Munich: P3 stocks. Complete genomic coverage was obtained for both the Wash1: P6 and Munich: P3 cDNA libraries. The Wash1 library was analyzed for 17 hours 27 minutes, generated 180.58 K reads and 79.95% of the passed reads mapped successfully to the genome sequence of Wash1. The Munich library was analyzed for 18 hours 2 minutes, generated 58.46 K reads and 56.53% of the passed reads mapped onto the Wash1 genome sequence. Both samples demonstrated increased coverage bias towards the 3’ end of the genome (Fig. 2A) which is commonly reported for cDNA libraries (34, 35). Basic variant detection was used to identify possible single and multiple nucleotide changes and deletions in the sequences encoding the S1/S2 region of the S protein (Fig. 2B). Twenty-two representative, single molecule reads obtained from the MinION sequencing of the Wash1: P6 stock over a 100 nucleotide region were selected, aligned (Fig. 2C), then translated and aligned (Fig. 2D) to published nucleotide and protein sequences respectively. The minority of the sequences in the MinION swarm retained the six amino acid consensus sequence (RRARSV). The majority had single and/or multiple nucleotide changes and/or deletions which removed the entire furin cleavage signal. These changes would be predicted to render the resulting S proteins furin-uncleavable and the consequences for studies of COVID-19 pathogenesis in animal models of disease may be significant. The Wash1: P6 stock also contained one other high frequency nucleotide change, resulting in an amino acid change (T323A), that was present with a frequency of 25% (Table 3). We did not observe any deletions in the MinION sequencing for the Munich: P3 stock, although there were four high frequency (greater than 25%) nucleotide changes resulting in amino acid changes (L368P, D614G, S686G and T941P; Table 3).

**Table 3.** Summary of single nucleotide changes, with frequency of 25% or greater, in the spike proteins of Munich: P3 and Wash1: P6 virus stocks determined by Oxford Nanopore cDNA-PCR sequencing

|  | reference position | reference<sup>1</sup> | change | read count<sup>2</sup> | coverage<sup>3</sup> | frequency (%)<sup>4</sup> | protein effect |
| --- | --- | --- | --- | --- | --- | --- | --- |
| Munich: P3 | 21640 | T | A | 199 | 243 | 81.89 | noncoding |
|  | 22665 | T | C | 14 | 49 | 28.57 | L368P |
|  | 23403 | A | G | 100 | 162 | 61.73 | D614G |
|  | 23618 | A | G | 58 | 217 | 26.73 | S686G |
|  | 24383 | A | C | 45 | 124 | 36.29 | T941P |
|  | 25075 | C | T | 6 | 17 | 35.29 | noncoding |
| Wash1: P6 | 21640 | T | A | 1471 | 1952 | 75.36 | noncoding |
|  | 22529 | A | G | 174 | 696 | 25 | T323A |

<sup>1</sup> reference sequence is CDC/2019n-CoV/USA\_WA1/2020 (accession number: MN985325)
<sup>2</sup> number of reads containing the change
<sup>3</sup> total number of reads across the reference position
<sup>4</sup> frequency of the change

**Figure 2.**
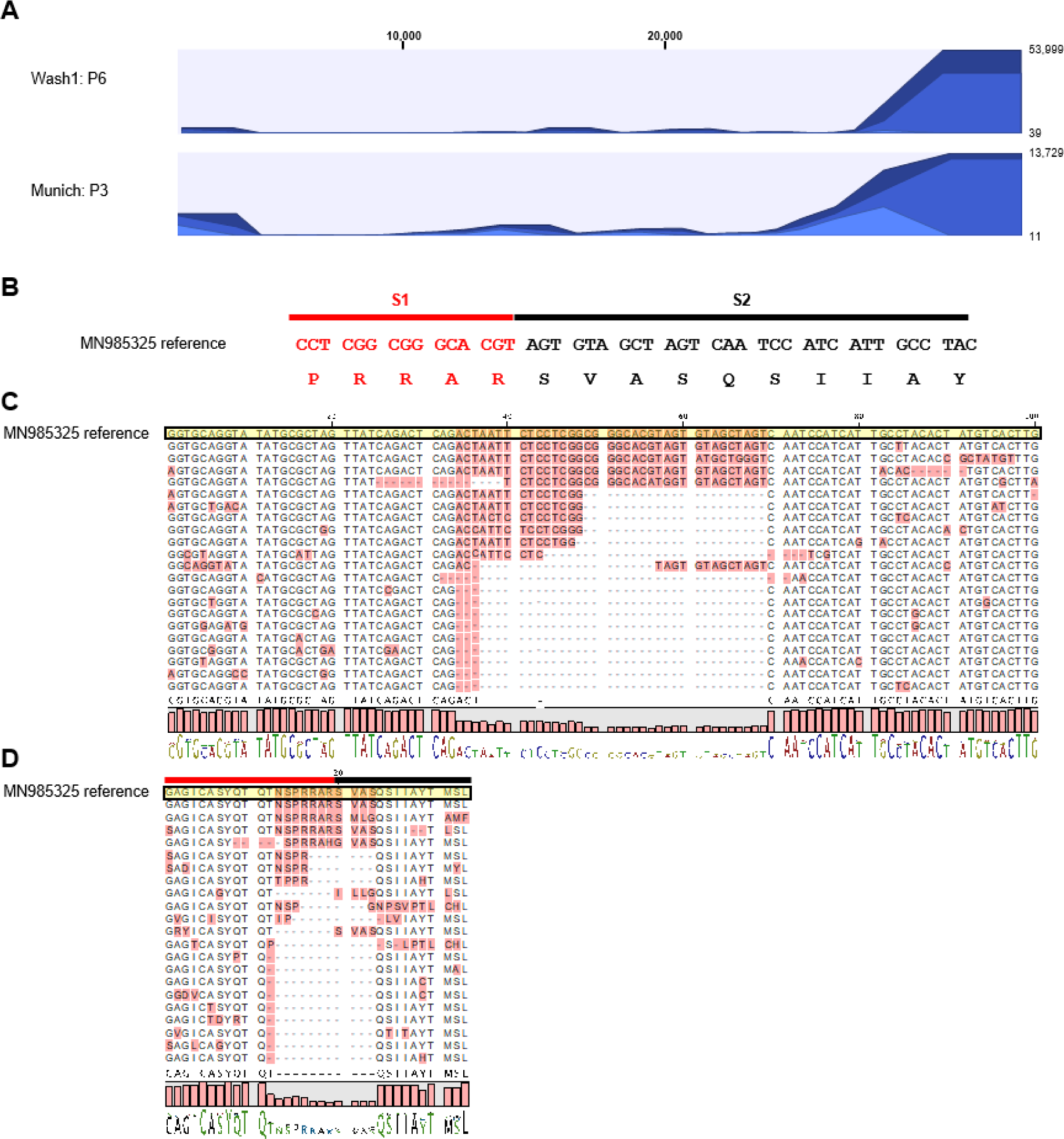
Virus adaptation. (A) Coverage of MinION sequencing reads. Full genome coverage was achieved for both Wash1: P6 (top) and Munich: P3 (bottom) virus stocks. Sequencing reads were aligned against the published Wash1 genome sequence (MN985325). The number of mapped reads are depicted on the y-axis with the nucleotide position within the SARS-CoV-2 assembly on the x-axis. (B) The S1/S2 region of the S protein including the furin cleavage signal (RRAR). Red and black lines indicate the putative S1 and S2 subunits, respectively, generated after cleavage. (C) Alignment showing representative, individual, mapped reads from the MinION sequencing to illustrate the diversity present in the nucleotide sequence coding for the furin cleavage signal region of the S protein. The reference sequence is shaded yellow and variable regions are shaded pink. The sequence logo at the bottom depicts the consensus sequence (letters) and it’s frequency (bar height). (D) Alignment showing the amino acid translations of the nucleotide sequences in (C) to illustrate the diversity present in the amino acid sequence of the furin cleavage signal (RRAR) region of the S protein. Red and black lines indicate the putative S1 and S2 subunits, respectively, generated after cleavage. The reference sequence is shaded yellow and variable regions are shaded pink. The sequence logo at the bottom depicts the consensus sequence (letters) and it’s frequency (bar height).

### Protein processing and localization

Since multiple deletions of the furin cleavage signal were present in the Wash1: P6 virus stock we analyzed cleavage of the S protein biochemically by developing an immunoblot to examine whole cell lysates prepared from virus-infected Vero-E6 cell monolayers (Fig. 3A). As expected the S protein from Wash1: P6 stock containing the deletions was uncleaved. However, the S protein was also uncleaved for a Wash1: P5 stock, in which the cleavage signal was intact (Fig. 1A). Similarly, the protein was not cleaved for either P2 or P3 Munich, for which the cleavage signal was intact (Fig. 1B). In contrast, an S protein expressed from a codon-optimized gene cloned into pCAGGS was cleaved efficiently (Fig. 3A, lane 2). An immunoblot performed using an antibody to the SARS-CoV-2 N protein on the same Vero-E6 cell lysates showed similar expression levels for each virus infection (Fig. 3A).

**Figure 3.**
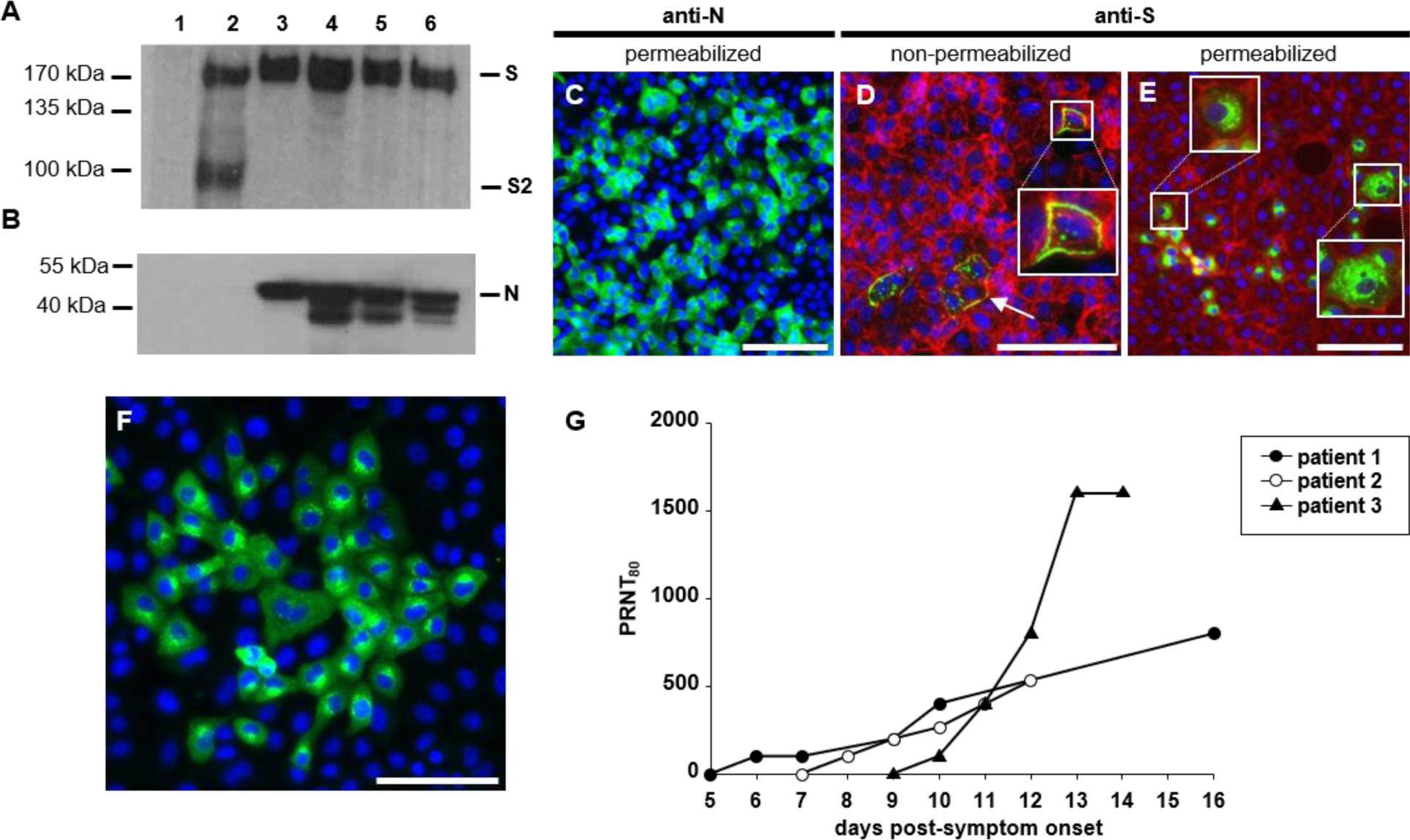
Antibody studies. (A) Western blot analysis of SARS-CoV-2-infected Vero-E6 cell lysates. Vero-E6 cells were infected with Wash1: P5 (lane 3) or P6 (lane 4) stock or Munich: P2 (lane 5) or P3 (lane 6) stock. Mock cell lysate prepared from cells transfected with a pCAGGS vector expressing eGFP (lane 1) and a lysate prepared from cells transfected with a pCAGGS vector expressing a codon-optimized S protein (lane 2) were included as a negative and positive control respectively. SARS-CoV-2 S protein and the S2 subunit, and the N protein, were detected with a SARS Coronavirus Spike protein polyclonal antibody or SARS Coronavirus N protein monoclonal antibody, respectively, and HRP-conjugated secondary. The predicted full-length S, S2 subunit and full-length N are indicated based on the marker sizes. Loading of cell lysates was adjusted to show equal levels of virus proteins. (B-D) Virus protein distribution in Munich: P3-infected Vero-E6 cells at 24 hours post infection visualized by immunodetection of N protein (B; green) or S protein (C and D; green) in permeabilized (B and D) or non-permeabilized (C) cells. Filamentous (f)-actin and/or nuclei were counterstained with phalloidin (red) and DAPI (blue), respectively. Areas marked by white boxes are shown at higher magnification in the insets (D and E). An arrow marks S protein distributed across the cell membrane of an infected cell (D). Scale bars represent 100 µm. (E) Virus protein distribution in permeabilized, Munich: P3-infected Vero-E6 cells at 24 hours post infection visualized by immunodetection with human serum (patient 3, day 14, Table 1; green). Nuclei were counterstained with DAPI (blue). The scale bar represents 100 µm. (F) Development of neutralizing antibodies against SARS-CoV-2 Munich: P3 virus stock in sequential human serum samples during the acute phase of COVID-19. Mean PRNT_80_ values are joined by lines.

We established an indirect immunofluorescence assay to determine the expression and localization of SARS-CoV-2 N and S proteins in infected cells. Based on our virological and sequence analyses showing the furin cleavage site was intact, Vero-E6 cells were infected with Munich: P3 at an MOI of 0.01 for 24 hours. The N protein was present in a predominantly diffuse cytoplasmic distribution with some small areas of perinuclear accumulation (Fig. 3C). This is consistent with previous observations (36). We also studied S protein trafficking to the cell surface using Vero E6 cell monolayers that were infected as above and were either fixed and permeabilized or left non-permeabilized. In non-permeabilized cells, the S protein was present at significant levels in puncta, which were distributed across the cell membranes of infected cells (Fig. 2D, arrow). This is consistent with the assembly and egress of SARS-CoV-2 through membranes between the endoplasmic reticulum and the Golgi intermediate compartment. A close association of S protein with the actin cytoskeleton was seen when phalloidin was used to stain filamentous (f)-actin (Fig. 3D and E, red). In this and other assays (not shown) cytopathic effect involved cell rounding and detachment from the dish and a degree of cell-to-cell fusion was observed. Cells were permeabilized and equivalent indirect immunofluorescence was performed to examine the intracellular location of the SARS-CoV-2 S protein. A punctate distribution throughout the infected cell cytoplasm and the cell surface was observed (Fig. 2E). The same infection conditions and indirect immunofluorescence assays were used to examine antibody reactivity in human serum obtained from acutely infected human COVID-19 patients (Fig 3F). In addition to perinuclear puncta, consistent with S protein localization, the antisera recognized SARS-CoV-2 proteins diffusely distributed throughout the cytoplasm. Cell rounding and increased staining were observed, consistent with cells in the later stages of the infection.

### Virus neutralization with human sera

Given the readily detectable anti-SARS-CoV-2 antibody reactivity in the human serum (Fig. 3F), we obtained a series of sera from three acutely infected COVID-19 patients to determine if these neutralized the virus (Table 1). Sera were collected from patients at multiple d.p.s.o. and a PRNT_80_ assay was developed using the Munich: P3 stock. This SARS-CoV-2 virus neutralization assay was performed in triplicate for all samples to examine the kinetics of antibody development in hospitalized patients at varying stages of the infection (Fig. 3G). For all patients the first serum sample (ranging from 4-9 days after first reported symptoms) failed to neutralize SARS-CoV-2. However, seroconversion occurred in all patients the subsequent day and all samples neutralized. Assays were routinely validated using a non-specific human serum collected before circulation of SARS-CoV-2 in Pittsburgh (data not shown). For patient 1, neutralizing activity was first detected at 6 d.p.s.o. and steadily increased until 16 d.p.s.o. Likewise, the neutralizing activity of samples obtained from patient 2 increased with similar kinetics following the first detectable signal 8 d.p.s.o. Although neutralizing activity was first detected at 10 d.p.s.o. in patient 3 there was a remarkable difference in the rate of development and strength of the neutralizing response compared to patients 1 and 2. This patient had the highest titer of the three, reaching a stable PRNT_80_ of 1600 both at 13 and 14 d.p.s.o. This was at the higher end of titers published by other groups using equivalently timed sera and comparable assay conditions (37-41). Interestingly for the d.p.s.o. where samples were available for either two or all three of the patients, the titers were quite similar. Since the Wash1: P6 virus stock had deletions in the furin cleavage signal we investigated whether using this virus would confound a PRNT_80_ assay using serum (day 14) from patient 3. Both viruses were equivalently neutralized by this serum reaching the same PRNT_80_ of 1600. It is possible that this result was influenced by the cell line used for stock preparation as both viruses exhibited similar S cleavage when grown on Vero-E6 cells (Fig 3A).

## Discussion

Virus provenance is critical in studies that aim to understand pathogenesis and test interventions such as antivirals, antibodies and vaccines. The emergence of SARS-CoV-2 and the rapid proliferation of studies must therefore be augmented by solid virological assays and a comprehensive understanding of the pathogen genetically and immunologically. Here we systematically characterized European and North American isolates of SARS-CoV-2 passaged in Vero-E6 cells, as the majority of researchers use them worldwide.

Since we are studying SARS-CoV-2 pathogenesis in a wide range of animal species (including non-human primates [Hartman *et al*., in preparation], ferrets [Reed *et al*., in preparation], hamsters and mice) we generated large volume working stocks for the Wash1 and Munich viruses to ensure comparability and consistency across *in vivo* challenge experiments. Viruses were passaged twice in Vero-E6 cells and Sanger and MinION long read sequencing were used to characterize Wash1: P6 and Munich: P3 stocks. We were surprised to discover that the Wash1: P6 stock contained deletions in the sequence coding for the furin cleavage signal of the S protein in the majority of the MinION reads. Sanger sequencing confirmed the deletion in the P6 stock. However, there was no trace of the deletion in the chromatograms for the received P4 virus or a P5 stock, indicating that once a deletion occurs it is rapidly selected in Vero cells and enriched in the virus population. Other groups have reported similar perturbations of this region (36, 42-44), and the deletions have been linked to the development of a large plaque phenotype (36). We observed variations in plaque phenotype, although in our experience, plaque size and morphology is governed by the concentration of agarose or Avicell in the semi-solid overlay medium and varies significantly depending on the protocol used. We did however observe an increase in virus titer for Wash1: P6 stock, increasing from 2×10^5^ P.F.U./ml at P5 to 3.55×10^6^ P.F.U./ml by P6. The Munich strain, with no detectable cleavage site deletions, had titers of 5×10^5^ P.F.U./ml at P2 and 3.25×10^5^ P.F.U./ml at P3. Enhanced growth and higher titers are synonymous with cell-culture adaptation.

A study using retroviral pseudoviruses incorporating the SARS-CoV-2 S protein into virions (45) demonstrated that a mutant S protein in which the S1/S2 cleavage site region Q_677_TNSPRRAR/SV_687_ was changed to Q_677_TILR/SV_683_, removing the furin cleavage signal, facilitated enhanced entry of the pseudovirus into Vero-E6 cells. A further study using rhabdoviral pseudoviruses (24) concluded that TMPRSS2 activity is required for SARS-CoV-2 entry into the human lung cell line Calu-3 and a second study from the same group (46) proposed that furin mediated cleavage at the S1/S2 site promotes subsequent TMPRSS2-dependent entry into target cells. Vero cells are TMPRSS2-deficient (47), and SARS-CoV-2 entry into them is likely cathepsin-mediated (48). Taken together these studies suggest that there may be no advantage to retention of the furin cleavage signal in Vero cells and that presence of the sequence may actually be detrimental to entry of the virus into these cells. This explains why the deleted virus quickly became predominant in our studies. We did not detect a similar perturbation of the furin cleavage signal in the Munich stock, although this virus has had fewer total passages in Vero-E6 cells, and we postulate that once a similar change occurs it will be rapidly selected for with continued passage in Vero-E6 cells. The Munich strain of SARS-CoV-2, which was passaged three times less in Vero-E6 cells, had a more consistent genome copies-to-P.F.U. ratio and a stable overall titer as determined by plaque assay. From these data, we conclude that an increase of ∼1 log in SARS-CoV-2 titers should be considered indicative of cell adaptation.

Four high frequency (> 25%), single, non-synonymous, nucleotide changes *versus* the SARS-CoV-2 Wash1 genome sequence were detected in the Munich S gene (Table 3). D614G (49, 50) was present in all passages sequenced and has been suggested to increase the transmission efficiency of the virus (51). It has also recently been reported that the mutation allowed retrovirus pseudotypes to enter hACE2-expressing cells with greater efficiency (52). S686G, was first detected in P2 sequencing and was enriched upon passage in Vero-E6 cells. This change is located at the putative S1/S2 cleavage site with the amino acid, presumably, constituting the new amino-terminus of S2 upon cleavage. This serine residue, along with S673 and T678, has also been predicted to be subject to O-linked glycosylation, due to the inserted (relative to other members of the betacoronaviruses) proline at position 681 just upstream (53). It has been shown for Sindbis virus that when a mutation, introducing an N-linked glycosylation site, is placed adjacent to the furin cleavage site of the PE2 glycoprotein that cleavage is blocked (54). Whether the S686G change in Munich: P3 has any impact on cleavage efficiency and whether this site is in fact glycosylated, and the biological consequences thereof, remain to be determined experimentally. T941P is also a potentially significant mutation since it is located in the fusion core of heptad repeat 1, and modelling of changes in this region have recently been shown to lead to the loss of interactions stabilizing the post-fusion assembly (55).

An important observation was that S proteins containing intact furin cleavage signals remained uncleaved in infected Vero-E6 cell lysates. Although some studies report western blot analysis showing cleavage of the S protein (22, 24, 45, 46, 56), they examine S protein from purified pseudovirus and not SARS-CoV-2-infected Vero-E6 cell lysates. One study using human serum to detect proteins in infected Vero-E6 cell lysates (38) showed mainly uncleaved S protein with a trace of bands for S1 and S2 visible at the lowest serum dilution tested. Although it is possible the antibodies in the human serum predominantly recognize uncleaved S protein, it is likely that in this study the S protein was also not cleaved efficiently in the Vero-E6 cells. Our data suggest that, specifically in Vero-E6 cells, the S protein is not cleaved efficiently during protein biosynthesis in the infected cell but rather at a later stage, i.e. during packaging and release or upon attachment or entry to the next host cell. Follow up studies examining the cleavage state of the SARS-CoV-2 S protein on purified virions would be informative in this respect. Interestingly, the fact that the pCAGGS-expressed protein was cleaved efficiently in the cells, suggests either that the codon optimization of the S gene affects an aspect of protein expression or that the fully replicating SARS-CoV-2 suppresses cleavage in infected Vero-E6 cells. This is potentially important for vaccine developers who are using codon optimization.

Genome copies-to-P.F.U. ratios decreased for the Wash1 virus upon passage from P4-P6. Although we did not examine them, it is likely that this represents a reduction in virus particles. This potential reduction in virus particles may have implications for SARS-CoV-2 pathogenesis studies since it has been well documented with alphaviruses, also enveloped, positive-sense RNA viruses, that decreases in particle-to-P.F.U. ratios observed with passage correlate with enhanced infection efficiency for cells on which the viruses were amplified. With alphaviruses, this often involves accumulation of neutral or negatively charged to positively charged amino acid substitution mutations that enhance binding to cell surface heparan sulfate (33, 57-59). Furthermore, passaged viruses are generally attenuated in animal models of disease either through effects of efficient heparan sulfate binding on virus dissemination or reduction in actual particle doses given when plaque forming units were used to determine inocula (57, 58, 60, 61). However, we did not observe mutation to positively charged amino acids in the SARS-CoV-2 S protein with passage, suggesting that a different selective pressure might be in play.

The neutralization assays were developed using Munich: P3 stock since it had no deletions in the furin cleavage signal. Since the neutralizing antibody titers we determined are at the high end of those reported by other groups (37-41) we investigated whether there was any difference in neutralizing capacity against the Wash1: P6 mixed population virus stock, with a deletion in the cleavage signal. There was no difference in neutralization capacity between the two virus stocks. This suggests that the use of a virus containing the furin cleavage site deletion in such assays should not impact detection of neutralizing antibodies, or the titers thereof, at least when virus stocks are prepared on Vero E6 cells. Follow up studies, that lie beyond the scope of the present communication, using other disease-relevant cell types and cells over-expressing proteases such as furin, which may cleave the SARS-CoV-2 S proteins, will be critical to undertake as will the assessment of the relative levels of cleaved versus uncleaved S protein in infectious virus particles. These data demonstrating perturbation of the furin cleavage signal during cell culture adaptation led us to halt using Vero-E6 derived Wash1 stocks for pathogenesis studies in animals. An important conclusion from this work is that SARS-CoV-2 used to infect animals should be the lowest passage level possible and confirmed by sequencing. This will be especially relevant for efficacy studies of vaccines and antivirals.

## Authors and contributors

William B. Klimstra – conceptualization, methodology, investigation, formal analysis, supervision, writing – original draft preparation.

Natasha L. Tilston-Lunel – methodology, investigation, formal analysis, writing – review and editing.

Sham Nambulli – methodology, investigation, formal analysis, writing – review and editing.

James Boslett – methodology, investigation, formal analysis, writing – review and editing.

Cynthia M. McMillen – methodology, investigation, formal analysis, writing – review and editing.

Theron Gilliland – methodology, investigation, formal analysis, writing – review and editing.

Matthew D. Dunn – methodology, investigation, formal analysis, writing – review and editing.

Chengqun Sun – methodology, investigation, formal analysis, writing – review and editing.

Sarah E Wheeler – resources

Alan Wells – resources, writing – review and editing

Amy L. Hartman – conceptualization, methodology, investigation, formal analysis, supervision, writing – review and editing.

Anita K. McElroy – conceptualization, methodology, investigation, formal analysis, supervision, writing – review and editing.

Douglas S. Reed – conceptualization, methodology, investigation, formal analysis, supervision, writing – review and editing.

Linda J. Rennick – formal analysis, writing – original draft preparation.

W. Paul Duprex – conceptualization, methodology, investigation, formal analysis, supervision, writing – original draft preparation.

## Acknowledgements

We thank Dr. Natalie Thornburg (CDC Atlanta) and Drs. Christen Drosten and Jan Felix Drexler (Charité – Universitätsmedizin Berlin) for providing SARS-CoV-2 isolates (CDC/2019n- CoV/USA_WA1/2020 from the United States and Munchen-1.1/2020/929 from Germany). We appreciate the efforts of the University of Pittsburgh Department of Environmental Health and Safety for timely regulatory and logistical oversight of this work under pandemic conditions. We are grateful to Andrew Davidson and David Matthews for useful discussion regarding interpretation of MinION data. The help and encouragement of Kevin Washo, Chancellor’s Office, University of Pittsburgh, who actively supported these studies from the outside is gratefully acknowledged.

